# Depletion of all platelet integrins impacts hemostasis, thrombosis and tumor metastasis

**DOI:** 10.1101/2024.11.22.624871

**Authors:** Emily Janus-Bell, Cristina Liboni, Alexandra Yakusheva, Vincent Mittelheisser, Clarisse Mouriaux, Catherine Bourdon, Louis Bochler, Vincent Hyenne, Maria Garcia-Leon, Olivier Lefebvre, Jacky G. Goetz, Pierre H. Mangin

## Abstract

Platelet integrins, in addition to other platelet receptors, are known to control hemostasis, thrombosis but also metastatic progression. Yet, their exclusive but combined deficiency has never been tested in these processes. Taking advantage of PF4Cre-β1^−/−^/β3^−/−^ mouse strain, we show that platelets are exclusively depleted for all integrins. While they displayed impaired binding to fibrinogen and annexin-V, P-selectin exposure was normal. Platelet adhesion was abrogated on immobilized fibrinogen and fibrillar fibronectin under shear flow. PF4Cre-β1^−/−^ /β3^−/−^ mice presented an increased bleeding time and a profound defect in experimental models of arterial thrombosis. Platelet adhesion to tumor cells was also reduced, with a profound impact on tumor growth and metastatic burden in a model of triple negative breast cancer. Overall, these results confirm the central role of platelet integrins in hemostasis and thrombosis, and define their role in tumor growth and metastasis formation.

**40-word summary:** Depletion of all platelet integrins in PF4Cre-β1^−/−^/β3^−/−^ mice leads to increased bleeding time and inhibits *in vivo* arterial thrombosis. Integrin-null platelets reduce tumor growth and metastatic burden in orthotopic and experimental metastasis models. Platelet integrins control hemostasis, thrombosis and metastasis.

## INTRODUCTION

Platelets are small anucleated circulating blood cells, which play a central role in hemostasis. At site of vessel injury, they adhere, activate and aggregate to form a platelet plug that stops bleeding. Platelet adhesion at site of injury is molecularly initiated through GPIb-IX-V complex interaction with subendothelial von Willebrand factor (vWF) (Savage et al., 1996, 1998). GPIb-vWF bond formation is crucial for platelet attachment under elevated wall shear rates. Under lower shear conditions, platelet β1 and β3 integrins ensure the recruitment of circulating platelets by themselves (Savage et al., 1996, 1998). In addition, platelet integrins are particularly important for securing their stable adhesion to extracellular matrix (ECM) proteins. Once stably adhered, platelet activation is mostly initiated through GPVI – collagen interaction (Nieswandt et al., 2001; Nieswandt and Watson, 2003). This induces the conformational change of integrin αIIbβ3 towards a high affinity state for circulating fibrinogen (Takagi et al., 2002), inducing platelet aggregation and the formation of a platelet plug sealing the breach (Versteeg et al., 2013). Platelets are also key players in arterial thrombosis, which is initiated upon rupture of an atherosclerotic plaque in a diseased artery. This pathological process leads to the formation of a thrombus that results in life-threatening ischemic pathologies such as myocardial infarction or ischemic stroke (Jackson, 2011).

Integrins are heterodimer transmembrane receptors composed of an α and a β chain. Platelets express at their surface three β1 integrins, α2β1, α5β1 and α6β1 interacting with collagen, fibronectin and laminins, respectively (Beumer et al., 1994; Hindriks et al., 1992; Saelman et al., 1994; Sonnenberg et al., 1988); and two β3 integrins, αvβ3 and αIIbβ3 interacting with vitronectin and fibrinogen (Marguerie et al., 1979; Paul et al., 2003). The role of β1 integrins is limited to platelet adhesion and activation on ECM proteins, with no major role in hemostasis as evidence by a normal tail bleeding time in platelet α2, α5 and α6 deficient mice (Habart et al., 2013; Janus-Bell et al., 2021; Schaff et al., 2013). Furthermore, while no patient with a deficiency in one of the platelet β1 integrins has been identified, a patient with a depletion in α2 presented mild bleeding during childhood but then the risk of bleeding decreased in adulthood (Nieuwenhuis et al., 1986). Integrins α2β1 and α6β1, but not α5β1, are implicated in experimental thrombosis since their genetic depletion in mice lead to thrombosis reduction in several experimental models (He et al., 2003; Kuijpers et al., 2003; Schaff et al., 2013; Janus-Bell et al., 2021). Concerning platelet β3 integrins, the most abundant is αIIbβ3 bearing a central role in platelet aggregation through its interaction with circulating fibrinogen. αIIbβ3 role in hemostasis is evidence by Glanzmann thrombasthenia, an hemorrhagic disease caused by its absence or non-functionality (Solh et al., 2015). αIIbβ3 also plays a key role in arterial thrombosis as evidenced by currently used antithrombotic drugs targeting αIIbβ3 (van den Kerkhof et al., 2021).

Besides their key role in hemostasis and arterial thrombosis, platelets are also implicated in non-hemostatic functions such as embryonic and fetal development (Finney et al., 2012; Osada et al., 2012) or immunity (Tang et al., 2002; Yeaman, 2014). There is solid experimental evidence that platelets contribute to tumor metastasis: tumor cells (TCs) colonized much less efficiently the lungs in mice in the context of severe thrombocytopenia (Gasic et al., 1968). Since, the importance of platelets in tumor growth and metastasis has been confirmed in several experimental models (Nieswandt et al., 1999; Camerer et al., 2004; Coupland et al., 2012; Garcia-Leon et al., 2024; Mammadova-Bach et al., 2016). Platelets actively populate the tumor microenvironment where they modulate TCs proliferation, angiogenesis, immune cell infiltration and function (Li et al., 2024). Circulating platelets interact with TCs in the bloodstream, which protects them from shear forces (Egan et al., 2014) and immune attacks (Nieswandt et al., 1999) notably through inhibition of natural killer cells activity (Kopp et al., 2009) and the transfer of major histocompatibility complex class I to TCs (Placke et al., 2012). At distant sites, platelets favor TCs adhesion to the endothelium (Dardik et al., 1997; Pilch et al., 2002; Anvari et al., 2021; Garcia-Leon et al., 2024) and facilitate the extravasation process (Schumacher et al., 2013; Labelle et al., 2011; Garcia-Leon et al., 2024), which favors metastatic colonization. Post-extravasation metastatic foci have been shown to be populated with platelets where they further support metastatic outgrowth through immune modulation (Garcia-Leon et al., 2024). The action of platelets at distant sites is mediated by several receptors, including the GPIb-IX-V complex (Jain et al., 2007; Qi et al., 2018; Erpenbeck et al., 2010), CLEC-2 (Shirai et al., 2017; Ichikawa et al., 2020; Chang et al., 2015; Tsukiji et al., 2018; Sasaki et al., 2018) and GPVI (Jain et al., 2009; Mammadova-Bach et al., 2020; Garcia-Leon et al., 2024).

Platelet β1 and β3 integrins were reported to contribute to tumor metastasis. We previously reported that mice lacking all platelet β1 integrins developed less lung metastasis in both an experimental metastasis model and an orthotopic model, with α6β1 playing a primary role (Mammadova-Bach et al., 2016). Its implication could be dependent on its interaction with ADAM-9 expressed on TCs in the bloodstream (Mammadova-Bach et al., 2016), shielding them from shear stress and immune surveillance. Interestingly, the interaction of platelet α2β1 with MCF-7 breast cancer cells induces transforming growth factor β1 secretion, a cytokine known to promote epithelial-mesenchymal transition, likely suggesting this integrin to fuel in this way TCs invasive properties (Labelle et al., 2011; Zuo et al., 2019). In addition, αIIbβ3 has been reported to be implicated in tumor metastasis as αIIbβ3 inhibitors or its deficiency decrease experimental metastasis (Amirkhosravi et al., 1999, 2003; Bakewell et al., 2003; Honn et al., 1992; Zhang et al., 2012). In fact, αIIbβ3 is able to interact with TCs *via* bridging with fibrinogen or vWF (interacting on the other side with αvβ3 expressed on TCs surface), therefore contributing to the formation of a shield around TCs.

In this study, we characterize a mouse strain specifically deficient for all five platelet integrins (PF4Cre-β1^−/−^/β3^−/−^) which allows to distinguish integrin-based adhesion from other known receptors and to study the full contribution of platelet integrins by overcoming potential redundancies, that we know are important for platelet integrins (Janus-Bell et al., 2024). We ensured that depletion of integrins did not perturb expression levels of known platelet receptors and assessed the combined role of platelet integrins in the function on platelets, using flow cytometry and flow-based *in vitro* assays. Doing so, we confirmed the already known prominent contribution to hemostasis and arterial thrombosis using bleeding time assays and arterial thrombosis mouse models. We further demonstrate for the first time that they actively participate to tumor growth and confirmed their role in metastasis using orthotopic and experimental models of metastasis.

## MATERIALS AND METHODS

### Materials

U46619, PAR-4 peptide, Polydimethylsiloxane (PDMS) (Sylgrad 184 Silicone elastomer base and curing agent), cellular fibronectin, fatty acid free human serum albumin (HSA), Bovine Serum Albumin (BSA), Tween20, MgCl_2_, NH_4_Cl, KHCO_3_ and Citrate buffer pH 6.0, Antigen Retriever were from Sigma-Aldrich. Alexa Fluor 488-coupled fibrinogen (Cat. F-13191) and DIOC_6_ (3,3’-dihexyloxacarbocyanine iodide) were from Molecular Probes. Ethylenediaminetetraacetic acid (EDTA), iFluor 488-Phalloidin (1:1000), APC-Cy7-Viability Zombie NIR (1:1000), Annexin V-Alexa Fluor488 (used at 1 μg/mL) and eBioscience™ FOXP3/Transcription factor staining buffer set (Cat.00-5523-00) were from Invitrogen. Phosphate-buffered saline (PBS) was from ThermoFisher Scientific (DPBS 1X) and recombinant hirudin from Transgene. Alexa Fluor 488-annexin V was from Life Technologies (Cat. A13201) and paraformaldehyde (PFA) and Eukitt mounting medium (Cat.15320) were from Electron microscopy sciences. Adenosine diphosphate (ADP) was from Mast group. Convulxin was from Cryoprep. Fibrinogen was from Intertransfusion, vWF A3 binding peptide vA3-III-23 was from Cambcol and collagen was from Takeda (Kollagenreagens Horm). Ferric chloride (FeCl_3_) was from Prolabo. RPMI1640 and Fetal Bovine Serum (FBS) were from Dutscher. Penicillin/Streptomycin and puromycin were from Gibco. Luciferin was from Euromedex. Harris Hematoxylin Acidified (ref.6765003), Eosin Y alcoholic (ref.6766007) and Superfrost slides (Cat.J1800AMNZ) were from Epredia. Avidin/biotin blocking kit (Cat.SP-2001) was from Vector Laboratories and Fluoromount-G™ with DAPI (Fischer Scientific, Cat. 00-4959-52). Masson Trichrome kit, light green variation (Cat.361350-0000) was from CellaVision-RalDiagnostics, 12 well chambers (Cat. 81201) were from Ibidi and Tumor Dissociation kit-mouse (Cat. 130-096-730) from Miltenyi.

### Antibodies

For platelet glycoprotein expression: FITC-conjugated anti-αIIbβ3 (clone Leo.F2, Cat. M025-1, 1:20), anti-GPIbα (clone Xia.G7, Cat. M042-1, 1:20), anti-GPV (clone Gon.G6, Cat. M061-1, 1:20), anti-GPIX (clone Xia.B4, Cat. M051-1, 1:20) and anti-GPVI antibodies (clone Jaq.1, Cat. M011-1, 1:100) and anti-β3 (clone Luc.H11, Cat. M031-1, 1:20), were from Emfret Analytics. PE-conjugated anti-α2 (clone HMα2, Cat. 103506, used: 1 μg/mL), anti-α5 (clone 5H10-27, Cat. 103805, used: 2 μg/mL), anti-α6 (clone GOH3, Cat. 313612, used: 1 μg/mL), and Alexa Fluor 488-conjugated anti-β1 antibodies (clone HMβ1-1, Cat. 102211, used: 1 μg/mL), were from BioLegend.

For P-selectin exposure: FITC-coupled anti-P-selectin antibody (clone RB40.34, Cat. 561923, used: 2.5 μg/mL) was from BD Pharmingen.

For *ex vivo* tumor multiparametric flow cytometry: PE/Dazzle594 anti-CD103 (clone 2E7, Cat.121430, 1:100), PerCP/Cy5 anti-CD11b (clone M1/70, Cat. 101211, 1:500), PE anti-CD11c (clone N418, Cat. 117308, 1:200), APC anti-CD138 (clone, 281-2, Cat. 558626, 1:200), BV711 anit-CD19 (clone 6D5, Cat. 115554, 1:800), BV421 anti-CD25 (clone A18246A, Cat. 113705, 1:100), PerCP/Cy5 anti-CD3 (clone 17A2, Cat.100218, 1:50), Alexa Fluor 700 anti-CD4 (clone RM4-5, Cat. 100536, 1:1000), BV510 anti-CD45 (clone 30-F11, Cat.103138, 1:200), BV605 anti-CD8 (clone 53-6.7, Cat.100744, 1:200), PE anti-CTLA4/CD152 (clone UC10-4B9, Cat. 106306, 1:20), PE/Cy7 anti-F4/80 (clone BM8, Cat.123114, 1:100), PE/Cy7 anti-IgD (clone 11-26c.2a, Cat. 405720, 1:100), Alexa Fluor 488 anti-Ly6C (clone HK1.4, Cat. 128022, 1:1000), BV711 anti-Ly6G (clone 1A8, Cat. 127643, 1:200), Alexa Fluor 700 anti-MHCII (clone M5/114.15.2, Cat. 107622, 1:200), BV421 anti-NK1.1 (clone PK136, Cat. 108731, 1:100), PE/Dazzle594 anti-PD-1/CD279 (clone 29F.1A12, Cat. 135228, 1:200) and TruStain FcX™ (anti CD32/16 antibody, 1:50) were from BioLegend. PE anti-CD38 (clone 90/CD38, Cat. 553764, 1:300), PE anti-CD44 (clone IM7, Cat. 553134, 1:400), PE/Dazzle594 anti-CD80 (clone 16-10A1, Cat. 562504, 1:200), Alexa Fluor 647 anti-TCF-1 (clone S33-966, Cat. 566693, 1:300) were from BD Pharmingen. FITC anti-FoxP3 (clone FJK-16S, Cat. 53-5773-82, 1:50), PE/Cy7 anti-TIM-3/CD366 (clone RMT3-23, Cat. 25-5870-82, 1:200) were from eBioscience.

For tumor immunofluorescence: primary: anti-mouse CD45 biotinylated (clone#30-F11, Cat.103103, 1:100), secondary: Streptavidin-AF647 (Cat.405237, 1:200) were from BioLegend.

For immunofluorescence of platelets-TCs interactions: Alexa Fluor 647 coupled anti-RAM.1 (used 2 μg/mL) was produced in-house (EFS).

### Mice

We used C57BL/6J mice lacking platelet β1 and β3 integrins (PF4Cre-β1^−/−^/β3^−/−^) (Janus-Bell et al., 2024). PF4-Cre^+^ (called PF4Cre) mice served as controls. Male and female mice were used except for tumor models with murine triple negative breast cancer cell line where only females were used. All procedures for animal experiments were approved by the regional ethical committee for animal experimentation and the French government (animal facility authorization: G67-482-10 and APAFIS 27659-2020101308518816, C67-482-33 and APAFIS 37433-2022052016445806).

### Platelet count evaluation

After severing the tail of anesthetized mice, whole blood was collected into EDTA (6 mM). Platelet count was analyzed in an automatic cell counter (Element HT5, Heska).

### Platelet glycoprotein expression

After severing the tail of anesthetized mice, whole blood was collected into EDTA (6 mM). Blood was diluted with PBS to obtain 100,000 platelets/μL. Labeled murine antibodies were incubated with diluted blood for 30 min at room temperature. After incubation, blood was diluted with 500 μL of PBS and surface glycoprotein expression of 10,000 platelet events was determined by flow cytometry (Gallios, Beckman Coulter).

### Measurement of platelet soluble fibrinogen binding, P-selectin exposure and annexin V binding

Blood was drawn into hirudin anticoagulant (100 U/mL final concentration) from the abdominal aorta of anesthetized mice. For soluble fibrinogen binding and P-selectin exposure, whole blood was diluted into 0.35% tyrode albumin buffer containing hirudin (100 U/mL), incubated with 488-fibrinogen (20 μg/mL) or FITC-coupled anti-P-selectin antibody and stimulated with agonists (ADP 2 μmol/L, U46619 2 μmol/L or PAR-4 peptide 1 mmol/L) for 10 min. Samples were fixed by adding 50 μL of 4% PFA for 20 min and then diluted with 600 μL of PBS. For phosphatidylserine exposure, whole blood was diluted into 0.35% tyrode albumin buffer containing hirudin (100 U/mL), stimulated with agonists (convulxin 15 nmol/L or collagen-related peptide 1 μg/mL and PAR4 1 mmol/L) for 10 min and then incubated with Alexa Fluor 488-annexin V (1 μg/mL) for 20 min in the presence of hirudin. Fluorescence was determined on 10,000 platelet events by flow cytometry (Gallios, Beckman Coulter).

### *In vitro* flow based assay

Flow experiments were performed as previously described (Schaff et al., 2013). PDMS flow chambers (0.1 x 1 mm) were coated with fibrinogen (100 μg/mL), cellular fibronectin (300 μg/mL), vWF A3 binding peptide vA3-III-23 (100 μg/mL) or collagen (200 μg/mL) overnight at 4°C. Mechanical stretching of cellular fibronectin was performed by connecting 2 silicon tubes to vacuum pumps switched on at 100 mbar. Silicon tubes were clamp and, after 1 min, connected to the inlet and outlet of the flow chamber. The clamps were then opened to apply a tensile force to the surface and allow cellular fibronectin to multimerize through mechanical stretching. Flow channels were blocked with PBS containing HSA (10 mg/mL) for 30 min at room temperature to limit unspecific platelet adhesion. Hirudinated (100 U/mL) whole blood was drawn from the abdominal aorta of anesthetized mice and perfused through the channels with a programmable syringe pump (Harvard Apparatus, PHD 2000) at 300 s^-1^ (over fibrinogen, fibronectin and collagen) or at 1,500 s^-1^ (over vWF A3 binding peptide). Platelet adhesion or thrombus formation were monitored by differential interference contrast (DIC) (Leica DMI4000B) using a 63x or a 40x objective and a Hammamatsu ORCA Flash 4L.T camera (Hammatsu Photonics). The number of platelet adhering or the area of thrombus was quantified using Fiji.

### Bleeding time

Tail bleeding time was determined by severing 3 mm at the tip of the tail of anesthetized mice. The tail was immersed in 0.9% saline at 37°C and the time needed for the bleeding to stop was recorded. Tubes containing blood were spun for 5 min, the pellet were homogenized in lysis buffer (NH_4_Cl 150 mM, KHCO_3_ 1 mM, EDTA 0.1 mM, pH 7.2) and used for blood lost quantification as compared to a standard curve optical density at 540 nm. Standard curve was based on 50 μL of blood diluted in 450 μL of lysis buffer. Bleeding time was also determined by 25G needle puncture of the carotid artery of anesthetized mice. The time to cessation of bleeding after injury was determined using fluorescent macroscope (Leica Microsystems) coupled to a charge-coupled device (CCD) camera (ORCA-Fusion C14440-20UP, Hamamatsu). Image acquisition was performed with Metamorph software (Molecular Devices) and analysis with Fiji.

### *In vivo* thrombosis models

Mice received DIOC_6_ injection to label platelets. For the ferric chloride model, after exposition of the left common carotid artery, a vascular injury was induced by applying laterally to the carotid a 3×3 mm Whatman filter paper saturated with 7.5% FeCl_3_ for 2.5 min. For the pinching model, a mechanical injury was induced by pinching the abdominal aorta with forceps for 15 s. Thrombus formation was monitored for 20 min with a fluorescent macroscope (Leica Microsystems) coupled to a CCD camera (ORCA-Fusion C14440-20UP, Hamamatsu). Image acquisition was performed with Metamorph software (Molecular Devices) and analysis of the thrombus size with Fiji.

### Tumor cells

We decided to exploit a murine triple negative breast cancer cell line (AT3) which was previously established in the laboratory (Garcia-Leon et al., 2024). Cells were cultured at 37°C 5% CO_2_ in RPMI1640 supplemented with 10% Fetal Bovine Serum-FBS (Dutscher) and 1% Penicillin/Streptomycin under puromycin selection (1 μg/mL). At confluency, cells were trypsinized and sub-cultured for further passing.

### Platelets-tumor cells interaction (immunofluorescence)

For platelets-TCs interaction, 6,000 AT3-RedLuc-L2 (AT3) cells were seeded per well on a 12 well chamber and left to adhere overnight in complete medium. The following day, platelets (citrated platelet rich plasma prepared as in Garcia-Leon et al., 2024) were added to each well at the concentration of 500 platelets per cell and let to interact for 30 minutes at 37°C 5% CO_2_ in Tyrode’s buffer without MgCl_2_. After three PBS washes, cells were fixed in PFA 0.5% in PBS for 20 minutes at room temperature. After three PBS washes, cells were stained with Alexa Fluor 647 coupled anti-RAM.1 (2 μg/mL) and Phalloidin-iFluor 488 (1:1.000) for 1 hour at room temperature. Slides were then washed and mounted in Fluoromount-G™ with DAPI. Images were acquired with 60x/water objective at Olympus Spinning disk confocal microscope. Analysis was performed with Fiji. The number of cells interacting platelets was manually counted and data were then post-processed in Excel 6.0 and GraphPad Prism 9.0 for statistical analysis.

### Syngeneic orthotopic tumor model

Two-hundred fifty-thousand AT3 syngeneic triple negative breast cancer cells were intraductally injected in PF4Cre-β1^−/−^/β3^−/−^ and control (PF4Cre). Tumor dimensions were measured longitudinally up to 18 days post-injection (dpi) through Caliper and tumor volume was calculated as in the following formula (volume = width^2^ x length). At the sacrifice, tumors were excised, weighted and processed for *ex-vivo* flow-cytometry immunophenotyping or immunofluorescence/Masson staining. Together, lungs were harvested and processed for subsequent staining.

### Syngeneic experimental metastasis model

One-hundred-fifty-thousand AT3 cells syngeneic triple negative breast cancer cells were tail-vein injected in PF4Cre-β1^−/−^/β3^−/−^ and control (PF4Cre). Metastatic outgrowth was longitudinally followed *in vivo* by intraperitoneal injection of luciferin (5 μg/kg) and imaging with IVIS Lumina III (PerkinElmer) (Garcia-Leon et al., 2024). Fourteen dpi, mice were sacrificed. IVIS results were analysed with Living Image®, Excel 6.0 and GraphPad Prism 9.0 graphing the ratio of photon emission compared to D0 (Garcia-Leon et al., 2024).

### Mouse organs processing

Tumors and lungs were harvested and fixed in PFA 4%/PBS overnight and processed for paraffin inclusion as previously described (Garcia-Leon et al., 2024). Briefly, following two subsequent washes in PBS and ethanol 50%, tumors were dehydrated overnight in 70% ethanol. Later, tissues were included in an automated Leica inclusion machine according to this scheme: ethanol baths (2x at 70%, 1x at 80%, 1x at 95% and 2x at 100%) 1 hour each, 2 xylene baths (1 hour each) and 2 paraffin embedding baths (1 hour each).

### Hematoxylin and Eosin staining

Lung 10 μm slices were dewaxed and rehydrated with a decreasing alcohol scale until ethanol 70%. Slides were finally rinsed under a continuous stream of tap water for 5 minutes. Hematoxylin staining was carried out in Harris Hematoxylin Acidified for 3 minutes and then washed under a continuous stream of tap water for 5 minutes. Staining was differentiated with a quick wash in acid alcohol (1% HCl + 70% ethanol) and washed again under a continuous stream of water for 5 minutes. Counterstaining was carried out in Eosin for 15 seconds and washed again under a continuous stream of water for 5 minutes. Slides were then dehydrated in an increasing alcohol scale until a final bath in xylene and then mounted in Eukitt mounting medium. Images were acquired with Slide Scanner VS200 microscope (Olympus) with a 20x/air objective. Analysis was performed with QuPath 0.5.0 (Bankhead et al., 2017). Metastastic areas were manually annotated as the hematoxylin-dense areas. Total percentage of metastatic areas was calculated over the tissue area (automatically annotated *via* eosin signal thresholding).

### Immunofluorescence staining and analysis

Ten-μm slices from paraffin-embedded tumors were obtained with a Leica microtome on Superfrost slides. Slices were dewaxed and rehydrated with a decreasing alcohol scale and antigen was de-masked after boiling in Citrate buffer pH 6.0, Antigen Retriever (Garcia-Leon et al., 2024). Primary antibody (1:100) was incubated overnight at 4°C after background minimization in blocking solution (3% bovine serum albumin-BSA, 20 mM MgCl_2_, 0.3% Tween 20, 5% FBS in PBS) and endogenous avidin/biotin blockade. The following day, slides were incubated in Streptavidin-AF647 diluted 1:200 in BSA 5% (Sigma) for 1h at room temperature. Slides were mounted in Fluoromount-G™ with DAPI. Images were acquired with 40x/oil objective at Zeiss LSM800 confocal microscope. Analysis was performed with Fiji (Schindelin et al., 2012). Number of nuclei was automatically measured using “analyse particles” plug-in after background subtraction and thresholding on slides. The number of CD45^+^ positive cells was manually annotated on processed images. The percentage of CD45^+^ cells on the number of nuclei was expressed in Excel 6.0, while graph and statistical analysis were carried out in GraphPad Prism 9.0.

### Masson Trichrome staining and analysis

Paraffin-embedded tumors were cut in 5 μm slices with a Leica microtome on Superfrost slides. Following dewaxing in a degrading alcohol scale and hydratation, tissue slides were stained with Masson Trichrome kit, light green variation following manufacturer’s instruction modified with 5 min mordant and 10 min staining incubation. At the end, slides were dehydrated in a growing alcohol scale with two final baths in xylene and then mounted in Eukitt mounting medium. Images were acquired with Slide Scanner VS200 microscope (Olympus) with a 20x/air objective. Analysis was performed with QuPath 0.5.0 (Bankhead et al., 2017). Tissue area was thresholded and automatically annotated in the blue channel at high resolution (1.10 μm/px), while Masson positive area was thresholded and automatically annotated in the residual channel at very high resolution (0.55 μm/px). The analysis was manually corrected to remove tissue artifacts. Percentage of Masson positive area was calculated over the total tissue area in Excel 6.0 and statistical analysis was performed in GraphPad 9.0.

### *Ex-vivo* multiparametric flow-cytometry tumor immunophenotyping

A single cell suspension was obtained from freshly harvested tumors as in (Garcia-Leon et al., 2024). Briefly, 2 mm^3^ tumor pieces were cut in pieces before to be enzymatically digested using the Tumor Dissociation kit, mouse diluted in RPMI1640 according to manufacturer’s instruction. The digestion was carried out for 42 minutes at 37°C using the gentleMACS Octo Dissociator (Miltenyi, 130-096-427). Prior to antibody staining, the cell suspension was filtered through a 50 μm nylon mesh and red blood cells were lysed in ACK Buffer (150 mM NH_4_Cl, 10 mM KHCO_3_, 0.1 mM Na_2_EDTA). Gating of only viable cells was ensured with Viability Dye staining (Zombie NIR) for 15 minutes at room temperature in the dark, while antibodies’ aspecific binding was lowered thanks to the blockade of CD32/CD16 receptor for 20 minutes at 4°C. Next, cells were stained with primary conjugated antibodies (see antibodies section) for 15 minutes at 4°C. For intracellular staining, cellular suspensions were further fixed and permeabilized using the FOXP3/Transcription factor staining buffer set according to manufacturer’s instructions. Antibodies targeting intracellular antigens (FOXP3, TCF-1) were incubated for 15 minutes at 4°C. After washing, samples were acquired at Attune NxT (Invitrogen) flow cytometer and data were analyzed using FlowJo™ v10 Software (ThreeStar). Excel 6.0 and Prism GraphPad 9.0 were used for data post-processing.

### Statistical analyses

Statistical analyses were performed with GraphPad Prism software 9.5.0. Shapiro-Wilk normality test was applied before statistical test. Based on the Gaussian or non-Gaussian distribution of the data, unpaired t-test or Mann-Whitney test (in case of 2 groups comparison) was used. Welch correction (2 groups) was applied. In case of longitudinal analysis, Two-way ANOVA with Original FDR method of Benjamini and Hochberg was applied. Mantel-Cox test was applied for bleeding time. p- or q-values together with the number of replicates and independent experiments are indicated in figure legends.

## RESULTS AND DISCUSSION

### The central role of integrins in platelet function

To characterize the impact of all platelet integrins depletion, we took advantage of a mouse strain specifically deficient for platelet integrins (PF4Cre-β1^−/−^/β3^−/−^) (Janus-Bell et al., 2024). PF4Cre-β1^−/−^/β3^−/−^ mice are viable and fertile, but present reduced breeding, linked to an increased mortality during parturition. They have a 50% reduction in platelet count as compared to controls (PF4Cre: 10.7 x 10^5^ platelets/μL; PF4Cre-β1^−/−^/β3^−/−^ 4.58 x 10^5^ platelets/μL) (**Figure 1A**). The platelet surface expression of GPIbα, GPV, GPIX and GPVI was normal while, as expected, the levels of every integrin subunit was profoundly reduced (**Figure 1B**). Using flow cytometry, we observed that PF4Cre-β1^−/−^/β3^−/−^ platelets were unable to bind detectable levels of soluble fibrinogen in response to ADP or the PAR-4 peptide, due to the absence of αIIbβ3 (**Figure 1C**). In contrast, P-selectin exposure in response to the TxA2 analog U46619 or to the PAR-4 peptide, was completely normal, indicating no major role of platelet integrin in the secretion of the granule content **(Figure 1D**). Concerning annexin-V binding, we observed that it was markedly impaired in PF4Cre-β1^−/−^/β3^−/−^ mice compared to controls (**Figure 1E**). This result appears in contrast with the work of other groups demonstrating no impact or increase procoagulant platelet formation in Glanzmann patients (Aliotta et al., 2020), or with αIIbβ3 antagonists treatment (Hamilton et al., 2004) and it could result from the combined deficiency in all platelet integrins. Next, we evaluated the ability of integrins to support platelet adhesion to different surfaces under flow condition. Perfusing PF4Cre-β1^−/−^/β3^−/−^ mouse blood through microfluidic chips, we observed a complete inhibition of platelet adhesion over fibrinogen or fibrillar fibronectin at an arterial wall shear rate of 300 s^-1^ (fibrinogen, PF4Cre: 4.69 ± 0.96 x 10^4^ platelets/mm^2^; PF4Cre-β1^−/−^/β3^−/−^: 0.03 ± 0.02 x 10^4^ platelets/mm^2^; fibrillar fibronectin, PF4Cre: 1.30 ± 0.61 x 10^4^ platelets/mm^2^; PF4Cre-β1^−/−^/β3^−/−^: 0.07 ± 0.03 x 10^4^ platelets/mm^2^) (**Figure 1F-H**), indicating that β1 and β3 integrins are fundamental for platelets attachment and stable adhesion under relatively low shear flow. In sharp contrast, platelet adhesion to plasma vWF at 1,500 s^-1^ was normal (PF4Cre: 3.10 ± 0.24 x 10^4^ platelets/mm^2^; PF4Cre-β1^−/−^/β3^−/−^: 2.54 ± 0.18 x 10^4^ platelets/mm^2^) (**Figure 1I**), confirming the modest contribution of integrins to the transient adhesion of platelets to immobilized vWF under high shear rate (Savage et al., 1996). Overall, β1^−/−^/β3^−/−^ platelets are unable to bind to fibrinogen or fibrillar fibronectin under flow conditions.

**Figure 1:**
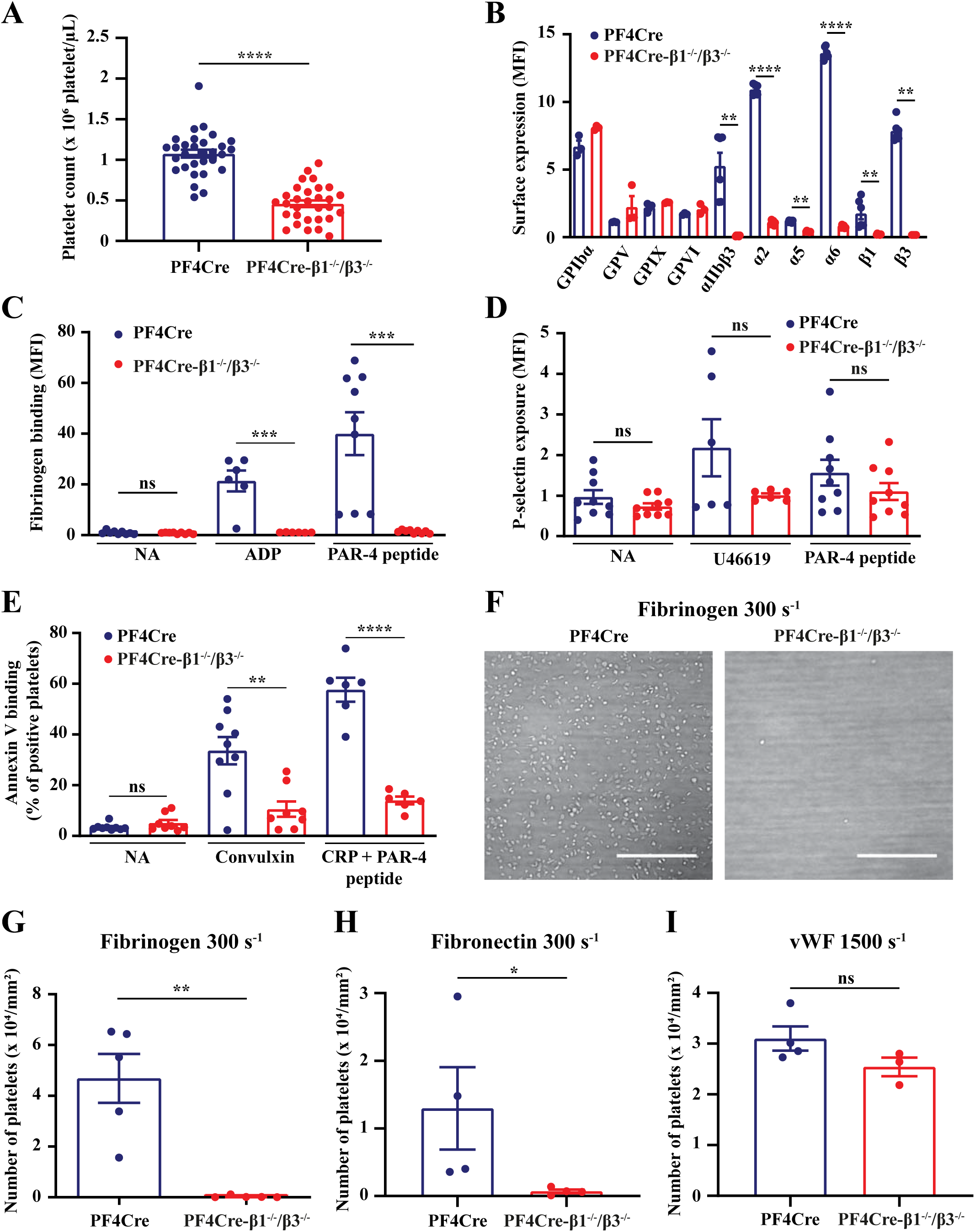
PF4Cre-β1^−/−^/β3^−/−^ mice platelet count, expression of major surface glycoproteins and platelet function. **A.** Platelet count. Platelet count from PF4Cre (n = 30) and PF4Cre-β1^−/−^/β3^−/−^ mice (n = 30) was evaluated. Data are from 6 independent experiments and unpaired t-test was applied. **B.** Glycoproteins surface expression. Surface expression of various glycoproteins on platelets in whole blood from PF4Cre (n = 3 or 6) and PF4Cre-β1^−/−^/β3^−/−^ mice (n = 3 or 6) was evaluated using selective antibodies and flow cytometry. Data are from 2 independent experiments and unpaired t-test or Mann-Whitney test were applied. **C.** Fibrinogen binding. Whole blood from PF4Cre (n = 6 or 9) and PF4Cre-β1^−/−^/β3^−/−^ mice (n = 6 or 9) was stimulated for 10 min with ADP (2 μmol/L) or protease-activated receptor 4 peptide (PAR-4 peptide) (1 mmol/L) and the binding of fluorescein isothiocyanate (FITC)-fibrinogen was detected by flow cytometry. Data are from 3 independent experiments and Mann-Whitney test was applied. **D**. P-selectin exposure. Whole blood from PF4Cre (n = 6 or 9) and PF4Cre-β1^−/−^/β3^−/−^ mice (n = 6 or 9) was stimulated for 10 min with U46619 (2 μmol/L) or PAR-4 peptide (1 mmol/L) and a FITC-conjugated anti-P-selectin antibody was detected by flow cytometry. Data are from 3 independent experiments and unpaired t-test or Mann-Whitney were applied. **E.** Annexin V binding. Whole blood from PF4Cre (n = 6 to 9) and PF4Cre-β1^−/−^/β3^−/−^ mice (n = 6 to 9) was stimulated with convulxin (15 nmol/L) or collagen related peptide (CRP) (1 μg/mL) + PAR-4 peptide (1 mmol/L) for 10 min, incubated with Alexa Fluor 488-annexin V for 20 min and analyzed by flow cytometry. The forward light scatter and fluorescence intensity of 10,000 cells were collected with a logarithmic gain and the percentage of annexin V-positive platelets was determined in the upper quadrant of the plot. Data are from 6 independent experiments and unpaired t-test or Mann-Whitney were applied. **F to I.** *In vitro* flow assays. Whole blood from PF4Cre (n = 4-5) and PF4Cre-β1^−/−^/β3^−/−^ mice (n = 3-5) was perfused through PDMS flow chambers coated with fibrinogen (100 μg/mL) at 300 s^-1^ for 5 min (F and G) or with fibrillar cellular fibronectin (300 μg/mL) at 300 s^-1^ for 8 min (H) or with vWF A3 binding peptide vA3-III-23 (100 μg/mL) at 1,500 s^-1^ for 2 min (I). Platelet adhesion was visualized in random fields by DIC microscopy, scale bar 50 μm (F), and the number of adherent platelets was quantified (G, H and I). Data are from 2 or 3 independent experiments and unpaired t-test (I) or Mann-Whitney (G and H) were applied. Results are expressed as the mean ± standard error of the mean (SEM); ns p > 0.05; * p < 0.05; ** p < 0.01; *** p < 0.001; **** p < 0.0001.

### PF4Cre-β1^−/−^/β3^−/−^ mice exhibit a major defect in hemostasis and experimental thrombosis

As the combined role of platelet integrins in thrombus formation was never characterized before, we assess their involvement in this process. Hence, anticoagulated whole blood of PF4Cre-β1^−/−^/β3^−/−^ mice was perfused through a microfluidic chip coated with immobilized type-I collagen at 300 s^-1^. Aggregation was abolished with PF4Cre-β1^−/−^/β3^−/−^ blood, where only single platelet adhesion to the surface was observed (area of thrombus, PF4Cre: 2.9 ± 1.0 x 10^4^ μm^2^; PF4Cre-β1^−/−^/β3^−/−^: 0.0 ± 0.0 x 10^4^ μm^2^) (**Figure 2A, B**). This result highlights the instrumental role of platelet integrins in thrombus formation under flow condition, but also questions the mechanism by which platelets are able to stably adhere over collagen in absence of α2β1, the most known platelet adhesion receptor to collagen. We speculate that the combined role of the GPIb-IX-V complex, binding to collagen-bound vWF, coupled to GPVI binding to collagen, might explains the residual PF4Cre-β1^−/−^/β3^−/−^ platelet attachment to the surface. Considering this major defect in thrombus formation, we then move to assess the bleeding time. Using a well-characterized mouse tail bleeding time model, we observed that PF4Cre-β1^−/−^/β3^−/−^ mice present a very marked increase in bleeding time, forcing us to stop the experiment at 600 s by cauterizing the injury to avoid the mice to bleed to death (**Figure 2C**). In addition, the volume of blood loss was markedly augmented in PF4Cre-β1^−/−^/β3^−/−^ mice as compared to controls (PF4Cre: 17.0 ± 9.74 μL; PF4Cre-β1^−/−^/β3^−/−^: 344 ± 34.3 μL) (**Figure 2D**). These findings were confirmed in a second hemostasis model consisting in a needle puncture of the carotid artery, which is much more severe than the classical tail-bleeding assay (**Figure 2E**). This increase in bleeding time is not explained by the 50% reduction in platelet count, as thrombocytopenic mice only present an increased bleeding time when the platelet count is below 100,000 platelet/μL (Morowski et al., 2013). These results confirmed the expected major role of platelet integrins in hemostasis in mice, especially the one of αIIbβ3 which is evidenced by the severe bleeding phenotype observed in Glanzmann thrombasthenia patients (Solh et al., 2015).

**Figure 2:**
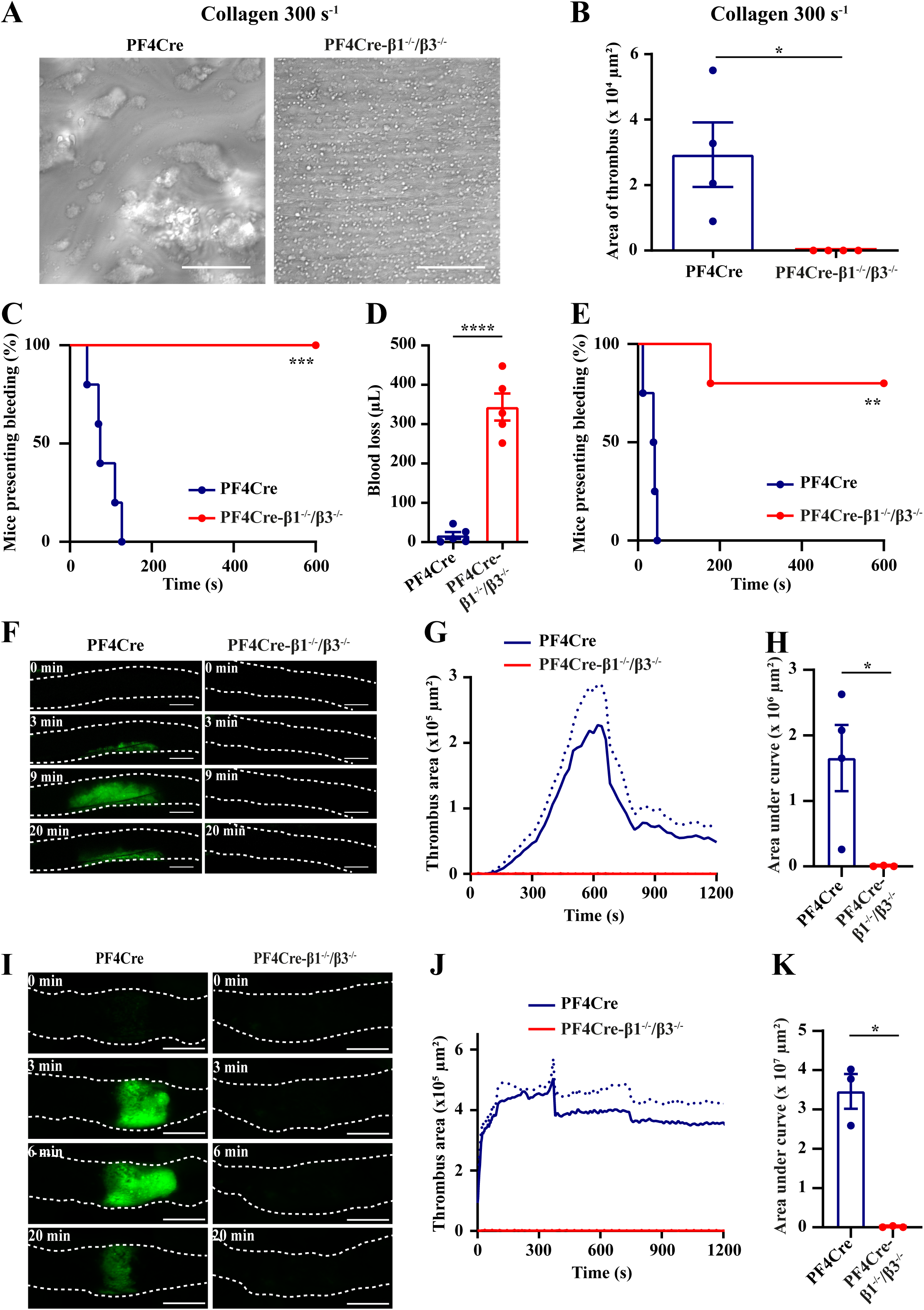
PF4Cre-β1^−/−^/β3^−/−^ mice are not able to form thrombosis and present an increased bleeding time. **A and B.** Thrombus formation *in vitro*. Whole blood from PF4Cre (n = 4) and PF4Cre-β1^−/−^/β3^−/−^ mice (n = 4) was perfused at 300 s^-1^ through PDMS flow chambers coated with collagen (200 μg/mL) for 8 min. Platelet aggregation was visualized in random fields by DIC microscopy, scale bar 50 μm (A). The area of the thrombi was quantified (B). Data are from 2 independent experiments and Mann-Whitney was applied. **C and D.** Tail bleeding time. The tail of PF4Cre (n = 5) and PF4Cre-β1^−/−^/β3^−/−^ mice (n = 5) was sectioned. The time required for the bleeding to stop was recorded (C) and the volume of blood lost (D) was measured. Data are from 1 experiment and Mantel-Cox test (C) or Mann-Whitney test (D) were applied. **E.** Carotid bleeding time. The carotid artery of PF4Cre (n = 4) and PF4Cre-β1^−/−^/β3^−/−^ mice (n = 4) was punctured and the time required for the bleeding to stop was measured. Data are from 1 experiment and Mantel-Cox test was applied. **F to H.** FeCl_3_ thrombosis of the carotid artery. Thrombosis was triggered in PF4Cre (n = 4) and PF4Cre-β1^−/−^/β3^−/−^ mice (n = 3) by application of a filter paper saturated with 7.5% FeCl_3_ to the common carotid artery. Representative fluorescence images of the thrombus (green) at the indicated time points after injury, scale bar 500 μm (F). The area of the thrombi was quantified (G) and area under the curves (AUC) are used to compare data (H). Data are from 1 experiment and Mann-Whitney test was applied. **I to K.** Pinching-induced thrombosis of the abdominal aorta. Thrombosis was triggered in PF4Cre (n = 3) and PF4Cre-β1^−/−^/β3^−/−^ mice (n = 3) by compression of the abdominal aorta with forceps. Representative fluorescence images of the thrombus (green) at the indicated time points after injury, scale bar 500 μm (I). The area of the thrombi was quantified (J) and AUC are used to compare data (K). Data are from 1 experiment and Mann-Whitney test was applied. The symbols correspond to individual mice. Results are expressed as the mean ± standard error of the mean (SEM); * p < 0.05; ** p < 0.01; *** p < 0.001; **** p < 0.0001.

As we only observed single platelet adhesion and no aggregation when perfusing PF4Cre-β1^−/−^/β3^−/−^ blood over collagen *in vitro*, we next evaluated the combined role of all platelet integrins in two mouse models of arterial thrombosis: a ferric chloride injury of the carotid artery and a mechanical injury of the abdominal aorta. We observed a very profound inhibition of *in vivo* thrombus formation in PF4Cre-β1^−/−^/β3^−/−^ mice with barely any sign of platelet adhesion at site of lesion (area under the curve for ferric chloride model, PF4Cre: 1.65 ± 0.51 x 10^6^ μm^2^; PF4Cre-β1^−/−^/β3^−/−^: 0.005 ± 0.005 x 10^6^ μm^2^; for mechanical injury: PF4Cre: 3.46 ± 0.44 x 10^7^ μm^2^; PF4Cre-β1^−/−^/β3^−/−^: 0.01 ± 0.01 x 10^7^ μm^2^) (**Figure 1F-K**), highlighting the instrumental role of mouse platelet integrins in thrombus formation also *in vivo*. Similarly to the bleeding time, these results are not explained by the 50% reduction in platelet count, as mice with a platelet count above 300,000 platelets/μL in a model of mechanical injury and 200,000 platelets/μL in a model of ferric-chloride injury, present normal experimental thrombosis (Morowski et al., 2013). In summary, these results demonstrate that integrins are central to hemostasis and arterial thrombosis as they finely control platelet adhesion and aggregation at the site of vascular injury.

### PF4Cre-β1^−/−^/β3^−/−^ mice present a reduced primary tumor and metastatic growth

We next aimed at testing whether combined integrin deficiency would alter platelet-dependent tumor phenotypes. Interestingly, adhesion of integrin-deficient platelets to triple-negative breast cancer cells (AT3 cells) was strongly reduced (PF4Cre-β1^−/−^/β3^−/−^: 0-3 platelets/cell; PF4Cre: 0-12 platelets/cell) **(Figure 3A)** prompting us to assess whether it could impact tumor progression. Platelets are a component of the tumor microenvironment and support primary tumor growth (Le Chapelain et al., 2024; Le Chapelain and Ho-Tin-Noé, 2022; Li et al., 2024). We first decided to investigate whether combined platelet β1 and β3 integrins deficiency might alter primary tumor growth using a syngeneic orthotopic model (AT3 cells) in PF4Cre-β1^−/−^/β3^−/−^ and PF4Cre mice (**Figure 3B**). Mammary tumor volume and weight were significantly reduced in PF4Cre-β1^−/−^/β3^−/−^ when compared to control mice (**Figure 3C**). Interestingly, lung metastasis were similarly perturbed in PF4Cre-β1^−/−^/β3^−/−^ mice, yet only mildly, suggesting that integrin deficiency is likely to impact multiple steps of metastatic progression (**Figure 3D**). Tumors of mice lacking platelet integrins had a tendency towards increased collagen deposition (**Figure 3E**). This suggests that absence of integrins in platelets impact their ability to organize the tumor-associated stroma, as seen in the light of the known role of platelets in ECM deposition (Malacrida et al., 2021) and integrin αIIbβ3 recognition as a driver of fibronectin fibrillogenesis *in vitro* (Lickert et al., 2022). In addition to these structural components, the stroma also comprises tumor-associated stromal cells including immune cells. Considering the well-established feedforward communication between platelets and the tumor immune microenvironment (Li et al., 2024), we further investigated this compartment by tumor immunofluorescence and observed a trend towards reduced infiltration of CD45^+^ cells in PF4Cre-β1^−/−^/β3^−/−^ derived tumors (**Figure 3F**). Additional analysis using *ex vivo* multiparametric flow-cytometry analysis of the tumor immune microenvironment initially confirmed that PF4Cre-β1^−/−^/β3^−/−^ tumors present a trend towards reduced number of CD45^+^ cells (**Figure 3G**). Within the myeloid panel, only neutrophils displayed an accumulation in PF4Cre-β1^−/−^/β3^−/−^ mice (**Figure S1**). With regards to the lymphoid lineage, while no variance in total T cells was observed, both CD4^+^ and CD8^+^ relative amount were slightly reduced in PF4Cre-β1^−/−^/β3^−/−^ tumors (**Figure 3G**). While a decrease in PD1-TIM3-compartment was observed both in CD4^+^ and in CD8^+^ cells (**Figure 3G**), PD1^+^ TIM3^-^ interferon producing effector/exhausted cells (Sakuishi et al., 2010; Fourcade et al., 2009) were exclusively increased in CD4^+^ subgroup (**Figure 3G**). In addition, T regulatory cells resulted increased in PF4Cre-β1^−/−^/β3^−/−^ mice (**Figure 3G**). While these data further question platelets’ role in immune cells recruitment at the tumor site (Schaubaecher et al., 2024), they demonstrate a non-negligible contribution of platelet integrins, as suggested by previous studies showing that platelets could promote anti-tumor T cells function *via* microparticle adoptive transfer of β3 integrin in a hepatocellular carcinoma model (Zhou et al., 2024). Alternatively, platelet integrins might alter the surface expression of immune checkpoint molecules hence remodeling T cell response (Schaubaecher et al., 2024). Overall, our data point towards a key involvement of combined platelet β1 and β3 integrins deficiency in impairing primary tumor growth, ECM deposition and immune cells recruitment and function.

**Figure 3.**
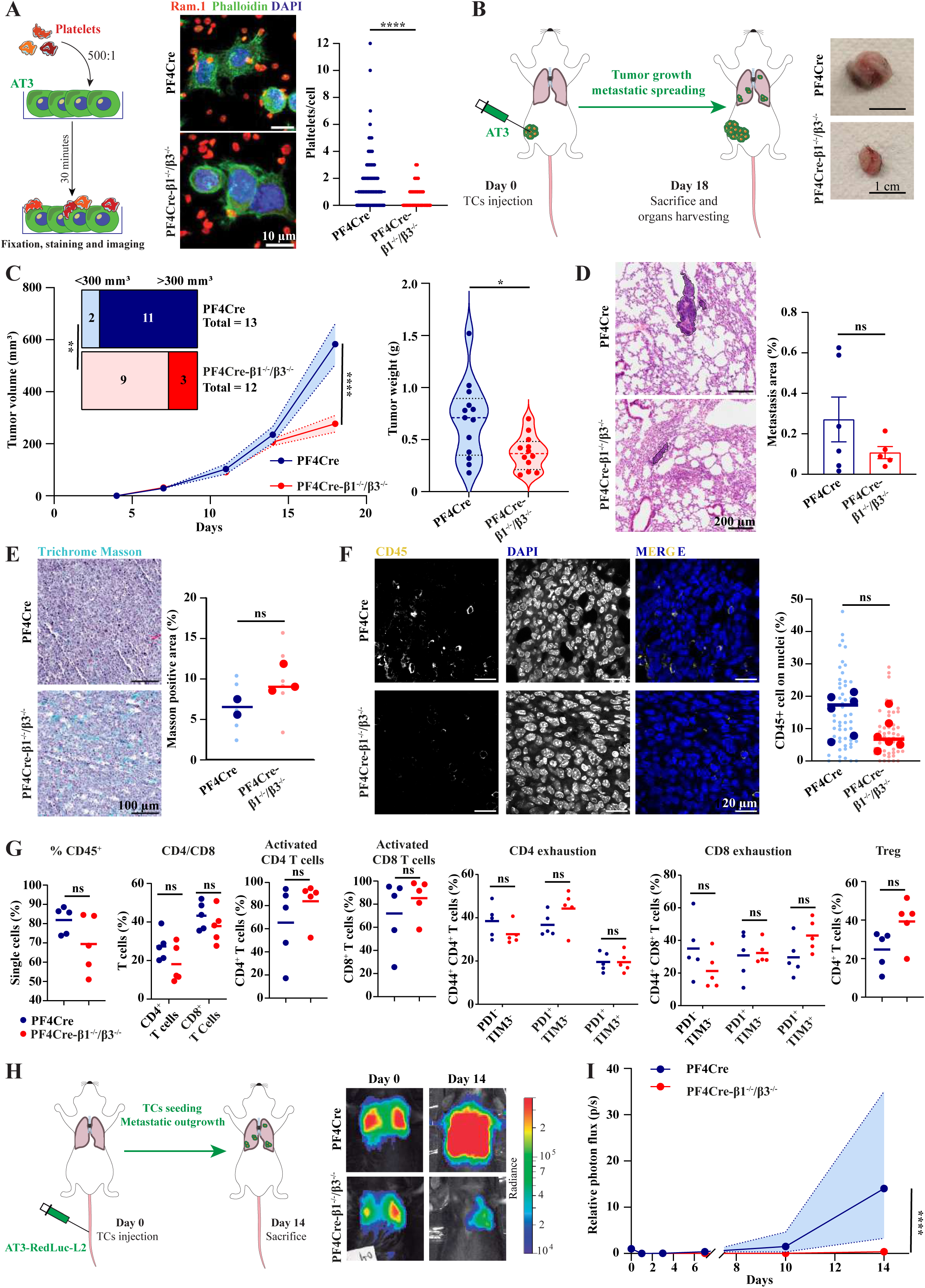
PF4Cre-β1^−/−^/β3^−/−^ platelets promotes a reduced binding to tumor cells, primary tumor growth and metastatic outgrowth. **A.** Platelets-Tumor cells *in vitro* interaction. Left: Experiment infographics. Middle: Representative confocal microscopy images (scale bar 10 μm). Right: graph representing the number of interacting platelets per single tumor cell for PF4Cre (356 cells) and PF4Cre-β1^−/−^/β3^−/−^ (288 cells). Data are from 1 experiment and Mann-Whitney test was applied. **B.** Orthotopic tumor model. Left: Infographics of the experimental scheme of the *in vivo* orthotopic tumor model. Right: Representative images of PF4Cre and PF4Cre-β1^−/−^/β3^−/−^ tumors at the sacrifice. Scale bar 1 cm. **C.** Orthotopic tumor model results. Left: Longitudinal tumor model measurement *in vivo* (measured *via* caliper) from PF4Cre (n = 13) and PF4Cre-β1^−/−^/β3^−/−^ mice (n = 12) during the course of the experiment. Mean ± range is represented (dotted lines). Data are from 2 independent experiments and Two-way ANOVA was applied. On the upper left, classification of number of mice with different tumor volume. The classification method is indicated in the figure. Right: *Ex-vivo* tumor weight measurement in PF4Cre (n=13) and PF4Cre-β1^−/−^/β3^−/−^ (n=12). Mean, quartiles and data distribution are represented. Data are from 2 independent experiments and Mann-Whitney test was applied. **D**. Hematoxylin and Eosin staining on paraffin-embedded lungs from orthotopic tumor model. Left: Representative images (scale bar 200 μm). Right: Quantification of the percentage of metastatic area on the total tissue area from PF4Cre (24 images from 6 mice) and PF4Cre-β1^−/−^/β3^−/−^ (20 images from 5 mice). Each dot represents a single mouse. Mean ± SD per experimental group is represented. Data are from 1 experiment. **E.** Masson Trichrome staining. Upper: representative images (scale bar 100 μm). Bottom: quantification of the percentage of Masson positive area from PF4Cre (6 images from 2 mice) and PF4Cre-β1^−/−^/β3^−/−^ (6 images from 3 mice). Each bigger point represents the mean value of a single mouse; littler points are representative of the mean of distinct images. Mean per experimental group is represented. Data are from 1 experiment and Mann-Whitney test was applied. **F.** Tumor Immunofluorescence on paraffin slides (IF) for CD45^+^ cells. Upper: representative images (scale bar 20 μm). Bottom: quantification of the percentage of CD45^+^ cells on the number of nuclei from PF4Cre (55 images from 6 mice) and PF4Cre-β1^−/−^/β3^−/−^ (52 images from 6 mice). Each bigger point represents the mean value of a single mice; littler points are representative of the mean of distinct images. Mean per experimental group is represented. Data are from 1 experiment and Mann-Whitney test was applied. **G.** *Ex-vivo* tumor immunophenotyping by flow-cytometry. Percentages of immune cell populations within the defined gating are represented for PF4Cre (n=5) and PF4Cre-β1^−/−^/β3^−/−^ (n=6). Data are from 1 experiment and Mann-Whitney test was applied. **H.** Left: Infographics of the experimental scheme of the *in vivo* experimental metastasis model. Right: Representative images at day 0 and day 14. **I.** Body Luminescence Index (BLI) from the experimental metastasis experiment. Relative photon flux (p/s) compared to day 0 is represented for PF4Cre (n=8) and PF4Cre-β1^−/−^/β3^−/−^ (n=6). Mean ± range is represented. Data are from 1 experiment and Two-way ANOVA was applied.

While understanding how platelets mechanistically control the recruitment of immune components requires additional investigation (Schaubaecher et al., 2024), they are known to bind to circulating TCs in the bloodstream where they foster metastasis (Labelle et al., 2014; Garcia-Leon et al., 2024). Since orthotopically-implanted tumors of PF4Cre-β1^−/−^/β3^−/−^ mice displayed a reduced, yet non-significant, metastatic potential (**Figure 3D**) and provide that adhesion of β1^−/−^/β3^−/−^ platelets to TCs was significantly reduced (**Figure 3A**), we set up an experimental metastasis model aiming at directly testing the contribution of platelet integrins to the hematogenous phase of metastasis (**Figure 3H**). As expected, metastatic outgrowth of AT3 cells was significantly reduced in PF4Cre-β1^−/−^/β3^−/−^ mice at 14 dpi (**Figure 3I**). This reduction in metastatic outgrowth is unlikely to be due to the 50% reduction of the platelet count as it has been described that 30% of the normal platelet count still allow a normal metastatic development (Coupland et al., 2012). While this result further recapitulates what was previously observed with platelet β1 deficiency (Mammadova-Bach et al., 2016) and β3 deficiency (Amirkhosravi et al., 1999; Bakewell et al., 2003; Honn et al., 1992; Zhang et al., 2012) or platelet integrins pharmacological blockade (Amirkhosravi et al., 1999, 2003; Bakewell et al., 2003; Honn et al., 1992; Zhang et al., 2012), it reinforces the central role of both platelet β1 and β3 integrins in mediating platelets-TCs interaction and subsequent metastatic progression.

In conclusion, this study characterizes PF4Cre-β1^−/−^/β3^−/−^ mice assessing the effect of all platelets integrins depletion on hemostasis, thrombosis and tumor growth. While integrins’ surface expression was completely abrogated, PF4Cre-β1^−/−^/β3^−/−^ platelets presented a normal glycoprotein surface expression and P-selectin exposure, indicating no major role of platelet integrins in granule content secretion. We provide evidence for a key role of integrins in many platelets’ function including platelet procoagulant activity and platelet attachment and stable adhesion at relatively low shear. In addition, aggregation over collagen under flow condition was completely blunted in the absence of platelet integrins, a notion which was confirmed *in vivo* by a complete prevention of thrombosis in two distinct models of arterial thrombosis. PF4Cre-β1^−/−^/β3^−/−^ mice presented severely increased bleeding in two distinct models of bleeding time, probably due to the absence of αIIbβ3, which reflects the severe phenotype observe in Glanzmann patients. Moreover, we showed that platelet integrins participate in platelets-TCs interaction. Their deficiency reduced primary tumor growth and, to some extent, pulmonary metastatic dissemination in a syngeneic AT3 model, impacting ECM deposition and immune cells accumulation and functionality at the tumor site. Furthermore, PF4Cre-β1^−/−^/β3^−/−^ mice presented diminished metastatic outgrowth in an AT3 experimental metastasis model. Overall, our study shed light on the irreplaceable functions of platelet integrins in dictating hemostasis and arterial thrombosis. In addition, we demonstrated their non-negligible participation in shaping primary and metastatic tumor growth.

## ACKNOWLEDGEMENTS

This work was supported by INSERM, EFS and ARMESA. We would like to acknowledge the PICSTRA imaging Platform for the technical support during imaging acquisition and Marina Peralta for Fiji analysis pipeline design. This work has been supported by funds from the French National Cancer Institute to P.M. and J.G.G. (PLBIO2023-173), which covers the financial support to C.L. V.M. is supported by a fellowship from the French Ministry of Science (MESRI) and a fourth-year thesis fellowship from the Fondation ARC pour la recherche sur le cancer. L.B. is supported by a PhD scholarship from FRM-Foundation pour la Recherche Medicale. Parts of the figures were created using material from Servier Medical Art. Servier Medical Art by Servier is licensed under a Creative Commons Attribution 3.0 Unported License (https://creativecommons.org/licenses/by/4.0/).

## AUTHORSHIP CONTRIBUTIONS

E.J.B. and C.L. acquired, analyzed, interpreted the data and wrote the manuscript; A.Y., V.M and O.L. acquired, analyzed, interpreted the data and contributed to the writing of the manuscript; C.M, C.B., L.B. and M.G.L. acquired, analyzed and interpreted the data; V.H. provided technical support; P.H.M. and J.G.G. conceived and designed the research, interpreted the data, wrote the manuscript and handled funding and supervision.

